# Errors associated with compound specific δ^15^N analysis of amino acids in preserved fish samples purified by high pressure liquid chromatography

**DOI:** 10.1101/812065

**Authors:** Rasmus Swalethorp, Lihini Aluwihare, Andrew R. Thompson, Mark D. Ohman, Michael R. Landry

## Abstract

Compound specific isotopic analysis of amino acids (CSIA-AA) is increasingly used in ecological and biogeochemical studies tracking the origin and fate of nitrogen (N). Its advantages include the potential for resolving finer-scale trophic dynamics than possible with standard bulk SIA and for reconstructing historical changes in the food webs of consumers from analyses of specimens in preserved sample archives. For the latter, assessing the effects of chemical preservatives on δ^15^N_AA_ has been inconclusive because the conventional CSIA approach for derivatized AAs by gas chromatography – combustion – isotope ratio mass spectrometry (GC-C-IRMS) has analytical errors (0.4 - 1.0 ‰) in the range expected from chemical preservation. Here, we show improved analytical precision (0.15 ± 0.08 ‰) for 11 underivatized AA standards analyzed by high pressure liquid chromatography followed by offline elemental analysis – IRMS (HPLC/EA-IRMS), an approach originally developed by Broek and McCarthy (2014). Using this method, we report the first high-precision tests of preservation effects on δ^15^N_AA_ in Northern Anchovy (*Engraulis mordax*) kept 1½-year in ethanol and up to 27-years in formaldehyde. We found minimal methodological induced fractionation for 8 AAs, and preservation effects on δ^15^N were similar regardless of duration and preservative used. Although some of the AAs differed significantly from frozen control samples (average +1.0 ± 0.8 ‰), changes in δ^15^N in the source AA Phenylalanine and trophic position estimates were statistically insignificant. Our results are encouraging for resolving fine-scale natural variability using HPLC/EA-IRMS on chemically preserved specimens and for ultimately reconstructing biogeochemical records and trophic dynamics over long time scales.

## Introduction

Compound specific analysis of nitrogen (N) stable isotopes in individual amino acids (CSIA-AA) is an increasingly common analytical method for tracking the origin and fate of N in ecological and biogeochemical studies. The method has proven particularly useful in disentangling food webs by identifying both the trophic positions of analyzed species and the base N sources utilized by primary producers (McMahon and McCarthy 2016; Ohkouchi et al. 2017). While bulk stable isotope analysis (bulk SIA) has been used to address these same ecological questions for many decades, interpretations are confounded by the integration of variances at both the trophic and source levels. CSIA-AA solves this problem by distinguishing the N isotope fractionation of specific AAs that enrich at a predictable rate with each trophic transfer, labeled “trophic” AAs, and those, labeled “source” AAs, that remain largely unaltered, reflecting inorganic source N at the food web base (Popp et al. 2007). By applying trophic discrimination factors (TDFs) derived from controlled feeding experiments or in situ comparisons of diet and isotopic fractionation patterns, it is possible to estimate the trophic positions (TP) of consumer organisms (McClelland and Montoya 2002; Chikaraishi et al. 2009; Bradley et al. 2015). Another advantage of CSIA-AA is that it provides baseline estimates that only consider the N sources actually consumed in the studied food chain, leading to more robust TP estimates (McClelland and Montoya 2002; Chikaraishi et al. 2007). Over the past decade, CSIA-AA has been used to resolve trophic connections within marine food webs and to track the flows of N and its inorganic sources (e.g., Chikaraishi et al. 2007; McCarthy et al. 2007; Hannides et al. 2009; Chikaraishi et al. 2010; Choy et al. 2012; Décima et al. 2013; Sherwood et al. 2014; Mompeán et al. 2016).

In trophic ecology studies, a major advantage of the CSIA-AA approach over bulk SIA is the acquisition of both trophic and source N isotopic measurements from samples of consumer tissue only. Thus, the extensive historical collections of various organisms in museum archives provide a potentially rich source of material for reconstructing past food webs and N cycling processes. Archived samples are in most cases chemically preserved, most commonly in formaldehyde or ethanol. Many studies have looked into the effects of such preservatives on bulk δ^15^N measurements and found variable and sometimes significant changes for a wide range of marine species (Rau et al. 2003; Kelly et al. 2006; Barrow et al. 2008). These occur mainly during the first few weeks to months of preservation (Sarakinos et al. 2002; Hetherington et al. 2019) and are likely the result of N compounds being solubilized and lost from preserved tissues since neither formaldehyde nor ethanol adds N (Bosley and Wainright 1999; Sarakinos et al. 2002). Much of an organism’s N is stored in AAs, but very little is known about the effects of chemical preservation on δ^15^N_AA_ fractionation or the impacts of multi-decadal storage. Few studies have explicitly tested the effects of preservation thus far. Hetherington et al. (2019) investigated short-term (2 year) effects of ethanol and formaldehyde in tuna and squid and the long-term (25 year) effects of formaldehyde on two species of copepods. Although the study did observe substantial but variable N isotopic fractionation, the preservation effects were not significant, in line with four other studies (Hannides et al. 2009; Ogawa et al. 2013; Strzepek et al. 2014; Chua et al. 2020).

One explanation as to why some studies have found statistically significant δ^15^N fractionation in chemically preserved bulk material but not in individual AAs may be uncertainties associated with the analytical methods used. Elemental Analyzer - Isotope Ratio Mass Spectrometry (EA-IRMS) used in bulk SIA typically operates with an analytical precision (0.1 - 0.2 ‰) that is substantially better than that reported for CSIA-AA (0.4 - 1.0 ‰; Bradley et al. 2016; Broek and McCarthy 2014; Broek et al. 2013; Chikaraishi et al. 2015; Hetherington et al. 2017; Nuche-Pascual et al. 2018; Ogawa et al. 2013; Ruiz-Cooley et al. 2017; Vane et al. 2018; Vokhshoori et al. 2019). In the few studies that have tested chemical preservation effects on δ^15^N_AA_, the degree of fractionation observed largely falls within the uncertainty range of the CSIA-AA approach (Hannides et al. 2009; Hetherington et al. 2019; Ogawa et al. 2013). The lower precision is the result of a lengthy and time-consuming analytical process inherent to the Gas Chromatography-Combustion-IRMS (GC-C-IRMS) method used for almost all CSIA-AA studies. For CSIA-AA, the AAs are first derivatized, causing significant and inconsistent fractionation of N isotopes, followed by a long and complex purification process prior to the δ^15^N measurement (Broek et al. 2013; Broek and McCarthy 2014). Ultimately, the propagation of errors limits the ability to detect and test the precise magnitude of chemical preservation effects. As an alternative to GC-C-IRMS, Broek et al. (2013) developed a method using High Pressure Liquid Chromatography (HPLC) for AA purification followed by offline EA-IRMS for δ^15^N measurement for Phenylalanine (Phe). Broek and McCarthy (2014) later optimized the method for both Glutamic acid (Glu) and Phe. Both studies compared results to GC-C-IRMS and found that, in addition to being a less expensive alternative, HPLC/EA-IRMS had better precision and accuracy for Glu and Phe.

To fully access and interpret the biogeochemical and ecological information locked away in historical archives, we first need to reduce analytical errors to a degree that will allow detection of the differences we expect to find. Only by doing so can we properly test if and to what extent isotopic signatures are modified by preservation methods, particularly during long-term storage. Minimizing error also has the advantage of increasing sensitivity and resolution of TP estimates. This is important for TP resolution for certain types of organisms, such as ammonium excreting bony fishes, which have TDFs of canonical trophic and source AAs approximately twice as large as the +3.4 ‰ δ^15^N trophic enrichment often used for bulk SIA studies (e.g., Zanden and Rasmussen 2001; Chikaraishi et al. 2009; Bradley et al. 2015). For this potential to be fully realized, however, CSIA-AA analytical errors must be comparable to those of bulk SIA.

In this study we first provide further evaluation of the HPLC/EA-IRMS approach to CSIA in underivatized AAs originally developed by Broek and McCarthy (2014). The evaluation is expanded from the original 2 AAs to 11 AAs of ecological interest, which more broadly resolves the analytical precision, accuracy and methodological reproducibility of HPLC/EA-IRMS. This allows us to assess the appropriateness of the method for testing the effects of chemical preservation relative to published errors from conventional GC-C-IRMS. We then use HPLC/EA-IRMS to test the effects of short-term (1½ year) preservation in ethanol and formaldehyde of white muscle tissue from adult Northern Anchovy (*Engraulis mordax*). In addition, we are the first to test the effects of long-term 27-year formaldehyde preservation of fish tissue (anchovy larvae) on δ^15^N_AA_ signatures. Based on former bulk SIA and CSIA-AA studies on marine fish, we hypothesize that the HPLC/EA-IRMS approach, being more sensitive than conventional GC-C-IRMS, will demonstrate significant δ^15^N fractionation as a result of chemical preservation. Lastly, we discuss the significances of methodology and chemical preservation on the precision and accuracy of TP estimation.

## Materials and procedures

### Amino acid standards

Tests of nitrogen isotope fractionation during sample processing, Amino Acid (AA) purification, and isotopic analysis were carried out using powdered standards and the liquid Pierce™ Amino Acid Standard H mix of 17 AAs. The powdered standards were L-Glutamic Acid (Glu), L-Alanine (Ala) and L-Proline (Pro) from Sigma-Aldrich, and L-Phenylalanine (Phe) from MP Biomedicals LLC. The 17 AAs in the Pierce standard mix were: Ala, L-Arginine (Arg), L-Aspartic Acid (Asp), L-Cystine (Cys), Glu, Glycine (Gly), L-Histidine (His), L-Isoleucine (Ile), L-Leucine (Leu), L-Lysine (Lys), L-Methionine (Met), Phe, Pro, L-Serine (Ser), L-Threonine (Thr), L-Tyrosine (Tyr), L-Valine (Val). Due to indications that our Phe powdered standard was contaminated (see results), we first purified 11 AAs of interest (Glu, Ala, Pro, Val, Ile, Leu, Phe, Gly, Ser, Tyr, Met) from the Pierce AA mix as a precaution by Liquid Chromatography (methodology described below). For both powdered and Pierce standards, half of each AA was taken as a control sample. The other halves were combined and split into four test samples. These test samples were not hydrolyzed, but were otherwise processed like fish samples and stored at −80°C for 1-4 weeks before AA re-purification (see below).

### Fish sample preparation

Adult Northern Anchovy (*Engraulis mordax*) were collected during the NOAA 2015 summer coast-wide coastal pelagic species survey off of central California (Zwolinski et al. 2016) using Nordic 264 rope trawl with a ∼600 m^2^ mouth and 8 mm mesh netting in the cod end liner that was towed at 3.5 knots, typically for 45 minutes. Within less than an hour of capture, two white muscle fillets were taken from the dorsal side of each individual fish (n = 6). One fillet was preserved in 95% tris-buffered ethanol and the other in 1.8% sodium borate-buffered formaldehyde. The remaining fish were stored at −20°C.

Larval anchovy were collected during the 1991 spring CalCOFI cruise from lines 80-83, stations 51-60 (www.calcofi.org) by oblique tows using Bongo nets of 0.71 m diameter, 0.505 mm mesh. The contents of one net were flash frozen in liquid N_2_ and stored at −80°C while the other side was preserved in seawater with formaldehyde (1.3% final concentration) buffered with sodium tetraborate. Larval anchovy of 8.5-10 mm in standard length (SL) were later sorted from the samples and stored at −80°C and in formaldehyde, respectively.

Adult anchovy samples for short-term preservation tests were defrosted or removed from the preservatives and a 16-51 mg dry weight (DW) sample of white muscle tissue taken for isotopic analysis. Preserved larval anchovy from long-term preservation tests were analyzed whole. For each test pair of frozen and formaldehyde-preserved samples, we ensured that the same proportion of larvae was taken from each 0.5 mm size interval and pooled 6-11 larvae to obtain enough material for isotopic analysis (0.5-2.0 mg DW).

All fish samples were frozen at −80°C and freeze-dried for 24 h. Adult tissue samples were then homogenized, and 0.5-0.8 mg subsamples taken for bulk isotopic analysis. The remaining homogenized adult tissue samples and the larval samples were stored in a desiccator until processing for CSIA-AA. A minimum of 400 µg of fish DW was hydrolyzed in 0.5 ml of 6N HCl in capped glass tubes for 24 hours at 90°C. Samples were then dried on a Labconco centrifugal evaporator under vacuum at 60°C, re-dissolved in 0.5 ml 0.1N HCl, and filtered through an IC Nillix – LG 0.2-µm hydrophilic PTFE filter to remove particulates. The samples were then re-dried before re-dissolving in 100 µl of 0.1% trifluoroacetic acid (TFA) in Milli-Q water, transferred to glass inserts in vials, and stored at −80°C for 1-4 weeks prior to AA purification.

### HPLC/EA-IRMS analysis of AAs and bulk material

The AA purification methodology is modified from the method of Broek and McCarthy (2014). We used an Agilent 1200 series High Pressure Liquid Chromatography system equipped with degasser (G1322A), quaternary pump (G1311A) and autosampler (G1367B). Samples were injected onto a reverse-phase semi-preparative scale column (Primesep A, 10 × 250 mm, 100 Å pore size, 5 μm particle size, SiELC Technologies Ltd.). Downstream a 5:1 Realtek fixed flow splitter directed the flow to an analytical fraction collector (G1364C) and an Evaporative Light Scattering Detector (385-ELSD, G4261A), respectively. We found that optimal AA detection on the ELSD was achieved with a nebulizer temperature of 30°C, evaporator tube temperature of 70°C, and a nitrogen gas flow rate of 1 L min^-1^ delivered from a nitrogen generator. A 120-min ramp solvent program with 0.1% TFA in Milli-Q water (aqueous phase) and HPLC grade acetonitrile (ACN, organic phase) was used as displayed on Fig. 1. Typically, the lifetime of the column was 200-250 runs before the chromatography deteriorated to a point where purification of Gly, Glu and/or Ala became compromised. A shorter program could be constructed, but required the system to operate at higher pressure (>250 bar), which translated into more maintenance from increased wear and tear. His, Lys and Arg can also be purified, but require a longer program (see Broek et al. 2013). A program was set up for the fraction collector to collect AAs of interest in 7 ml glass tubes at specific times based on elution times from previous runs. A steep gradient in δ^15^N can occur across the peak of an eluting AA (Hare et al. 1991; Broek et al. 2013). Therefore, the quality of all collections was assessed by comparing the chromatogram with set collection times, and only AAs where ≥ 99% of the peak areas fit within the collection windows were accepted. Due to slight drift in AA elution timing, collection times were modified between consecutive runs. Injection volume was determined from sample DW and expected content of low concentration AAs (typically Phe and Met) based on previous runs of similar samples. The aim was to collect approximately ≥1 µg N equivalent of each AA. Here, we injected samples of 484-776 µg DW of fish biomass, but it was possible to collect sufficient Phe for isotopic analysis from 350 µg DW.

**Figure 1:**
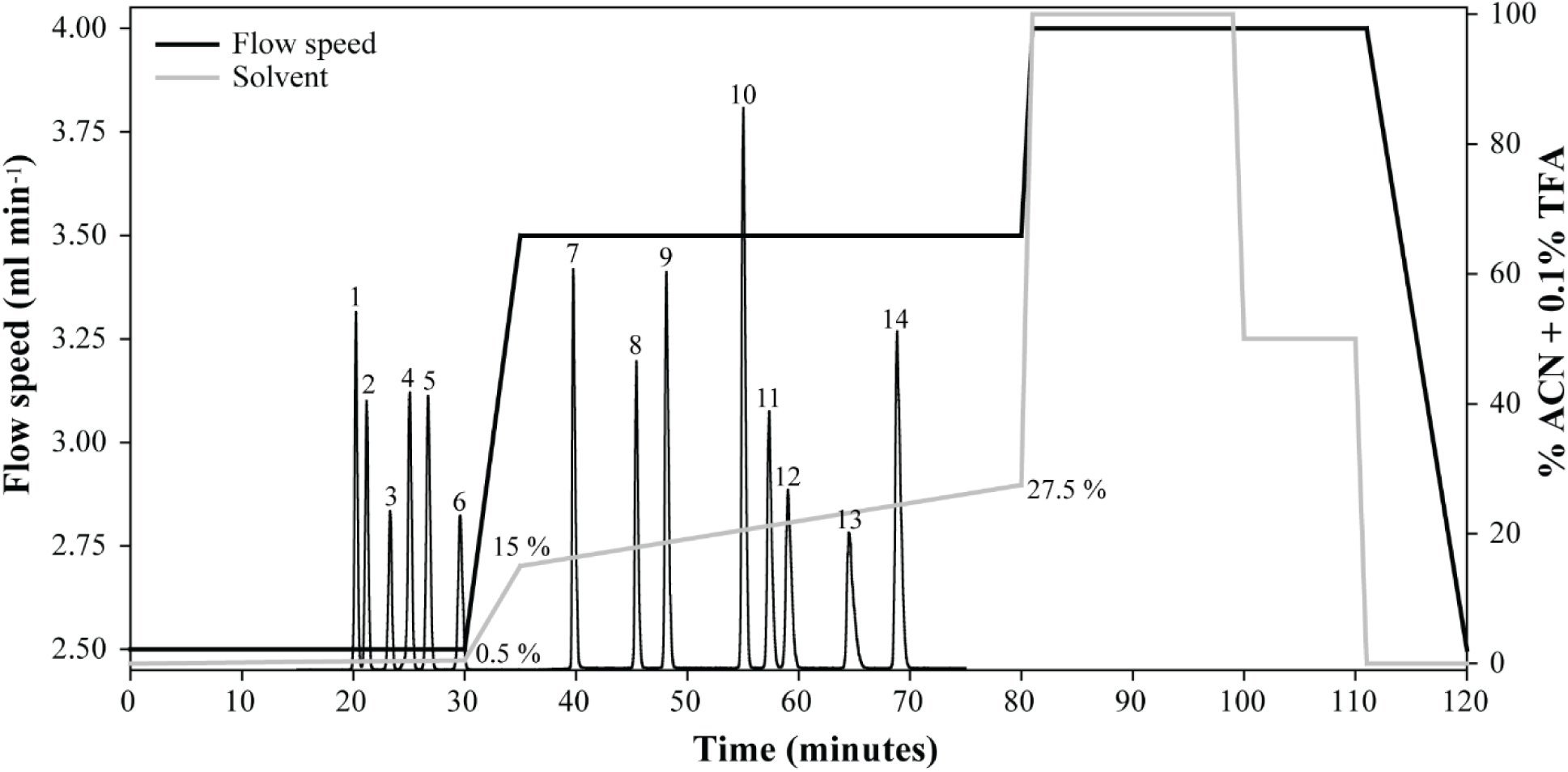
HPLC ramp solvent program modified from Broek and McCarthy (2014). Column cleaning and equilibration were carried out from time 80 to 120 min. System pressure typically ranged between 160 and 255 bars during a run. Background shows a chromatogram from injection of PierceTM AA mix. Peak identities are: 1. Asp, 2. Ser, 3. Gly, 4. Thr, 5. Glu, 6. Ala, 7. Pro, 8. Val, 9. Met, 10. Tyr, 11. Ile, 12. Leu, 13. Cys, 14. Phe.

Following collection, the AA samples were dried in the centrifugal evaporator at 60°C, dissolved in 40 µl of 0.1N HCl, and transferred to tin capsules (Costech, 3.5 x 5 mm). The capsules were then dried overnight in a desiccator under vacuum. Pre-combusted borosilicate glassware with PTFE lined caps was used for all process steps, and all sample transfers were done in HPLC-grade solvents or Milli-Q water with fine glass syringes.

Isotopic analysis of powdered AA standards and bulk material of white muscle tissue from adult anchovies was carried out at the Scripps Institution of Oceanography Stable Isotope Facility (SIO). Samples were analyzed on a Costech ECS 4010 Elemental Analyzer coupled to a Thermo Finnigan Delta Plus XP Isotope Ratio Mass Spectrometer. Sample ^15^N/^14^N ratios are reported using the δ notation relative to atmospheric nitrogen (N_2_). Measured δ^15^N values were corrected for size effects and instrument drift using acetanilide standards (Baker AO68-03, Lot A15467). Long-term analytical precision was on the order of ≤ 0.2 ‰.

Isotopic analyses of the Pierce AA standards and fish AAs were carried out at the Stable Isotope Laboratory facility at University of California, Santa Cruz (UCSC-SIL). Samples were analyzed on a Nano-EA-IRMS system designed for small sample sizes in the range of 0.8-20 µg N. The automated system is composed of a Carlo Erba CHNS-O EA1108 Elemental Analyzer connected to a Thermo Finnigan Delta Plus XP Isotope Ratio Mass Spectrometer via a Thermo Finnigan Gasbench II with a nitrogen trapping system similar to the configuration of Polissar et al. (2009). Measured δ^15^N values were corrected for size effects and instrument drift using Indiana University acetanilide, USGS41 Glu and Phe standards and correction protocols (see https://es.ucsc.edu/~silab) based on procedures outlined in Fry et al. (1992).

### Data analysis

TP was estimated using the equation, β, and trophic discrimination factor (TDF) values presented in Bradley et al. (2015) for individual trophic and source AA pairs. When calculating TP using multiple AAs, an average TP was taken of all trophic-source combinations. This was done to allow testing of TP estimates from individual AA trophic-source pairs.

Methodology effects on δ^15^N were tested by ANOVA followed by post hoc testing using Tukey HSD. Preservation effects on δ^15^N_AA_ and TP were tested by ANOVA followed by post hoc testing using paired two-way t-tests for each AA from frozen control and chemically preserved test material. Effects of preservation in bulk material were also evaluated by paired two-way t-tests. Before testing, data were inspected for normality of distribution followed by homogeneity of variance by the Levenes test. For the t-tests, we assumed unequal variance between control and test samples and p values adjusted using a Bonferroni correction. All statistical tests were carried out in R (R_Core_Team 2017).

## Assessment

### Methodological induced errors on individual AAs

Our methodological procedure for purifying individual amino acids (AA) did not cause significant fractionation of δ^15^N in most of the four target AAs (Fig. 2-a). One exception was Phe, which changed significantly relative to the control (+1.04 ‰, p < 0.001). This was also confirmed by a second test of Glu, Ala and Phe (results from both tests are pooled together in Fig. 2-a). For the other three AAs, a small but statistically insignificant fractionation was observed (−0.05 – 0.29 ‰, average: +0.12 ‰). This indicated that some of our standards might not have been sufficiently pure and prompted a third test for which we pre-purified 11 target AAs before testing to probably evaluate isotopic fractionation due to our processing and purification methodology. In this test for pre-purified standards, we found similar results to our initial tests (Fig. 2-b). Two exceptions were Ile and Met, which both changed significantly relative to control samples (−0.85 ‰, +2.11 ‰, p < 0.001). Although not significant, Tyr also showed substantial fractionation (+0.57 ‰, p = 0.059). For the other eight AAs, we found a high accuracy of +0.06 ‰ on average (Fig. 3). Glu, Ala and Phe showed virtually no fractionation (+0.01 - 0.04 ‰), while some, but statistically insignificant, fractionation was observed for Pro and Leu (+0.27 - 0.4 ‰, Fig. 2).

**Figure 2:**
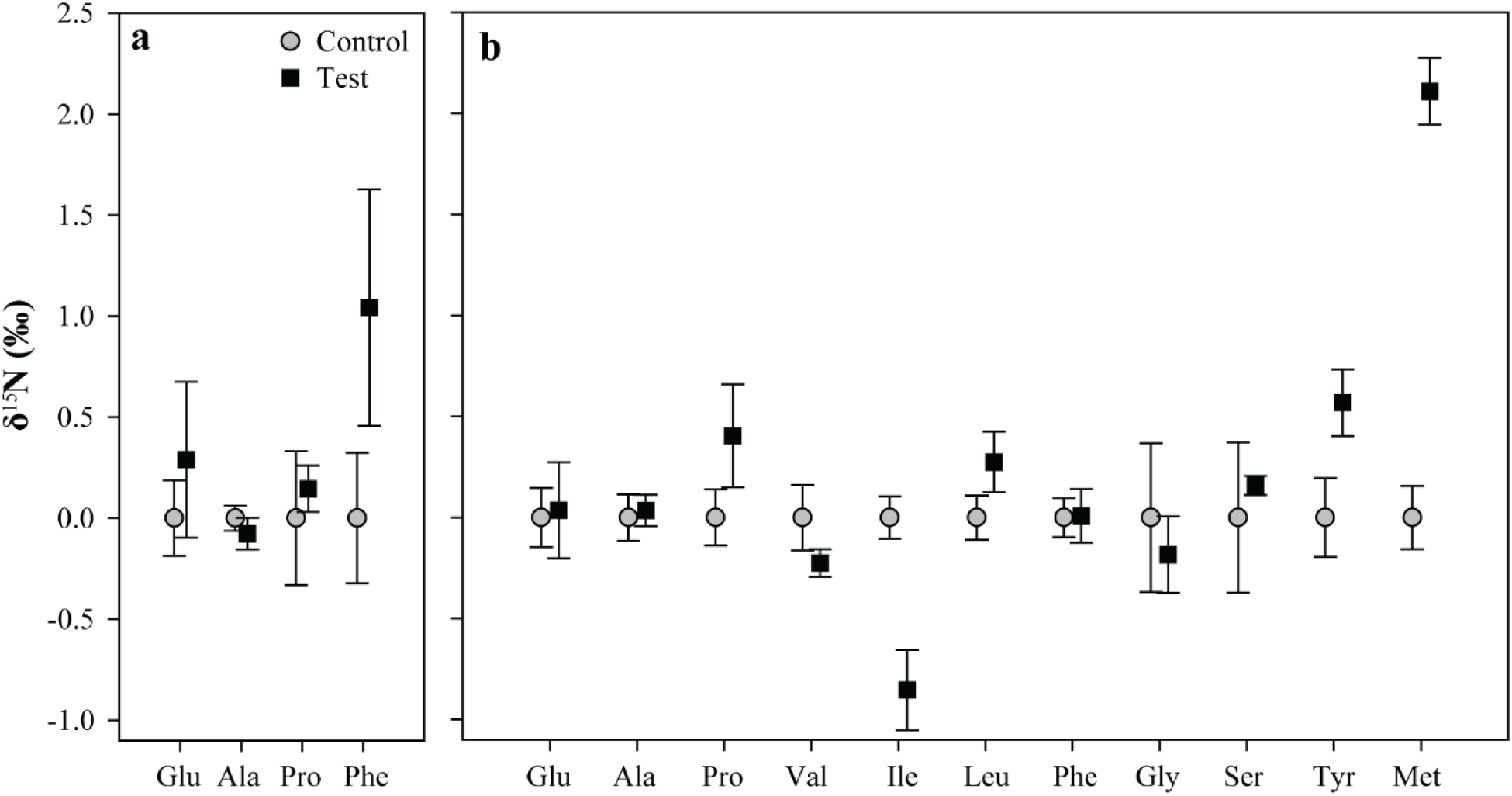
Change in δ15N in AA standards processed and purified by HPLC relative to control samples (±1 SD). A) shows results from two rounds of testing (except for Pro) using powdered standards analyzed on a “regular” EA-IRMS at SIO (Glu and Ala, n = 6; Pro, n = 2; Phe, n = 7), and B) shows results from testing using pre-purified standards analyzed on a Nano-EA-IRMS at UCSC (Glu, Phe and Gly, n = 4; Ala, Pro, Val, Ile, Leu, Ser, Tyr and Met, n = 3). n = minimum number of replicates per treatment.

**Figure 3:**
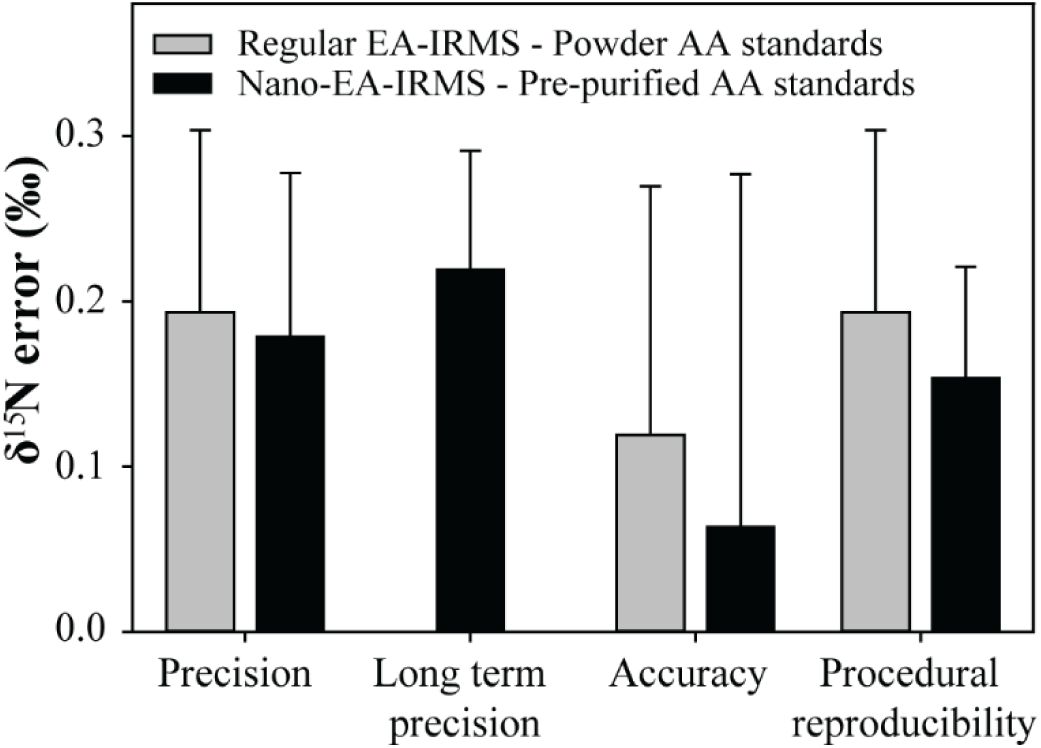
Errors associated with sample processing and analysis by HPLC/EA-IRMS. Analytical precision of the “regular” (n = 3) and Nano-EA-IRMS (n = 11) instruments at the SIO and UCSC isotope labs calculated as the average standard deviation for all AA control samples displayed in Figure 2. Long-term precision of Nano instrument is from 12 consecutive runs of powdered acetanilide and Phe standards over a one month period (n = 24). Procedural reproducibility is calculated as the average standard deviation for all AA test samples. Accuracy of the method is calculated as the average difference between test and control samples (excluding Ile, Tyr and Met, n = 8) displayed in Figure 2. The powdered Phe standard was not considered throughout this plot. Error bars are +1 SD.

The overall analytical precision of the Nano-EA-IRMS for all the AAs in Fig. 1 was within the range of the long-term performance of the instrument and similar to the precision of the regular EA-IRMS (Fig. 3). The accuracy between the powdered and pre-purified standards were similar, particularly when considering only Glu, Ala and Pro (+0.12 and +0.16 ‰, respectively). We also found the procedural reproducibility similar for tests using different standards (equal if considering only Glu, Ala and Pro), and comparable to the precision. Overall, the AA purification methodology did not add additional variability to the results.

### Preservation induced errors on individual AAs

Short-term (1½ year) preservation of adult anchovy white muscle fillets with ethanol and formaldehyde resulted in a significant fractionation of δ^15^N in bulk material (+0.67 ‰, t = 8.67, df = 5, p = 0.001; +0.84 ‰, t = 9.17, df = 5, p < 0.001; Fig. 4, Table S1). However, this fractionation was not consistent across all AAs or between preservative types and durations.

**Figure 4:**
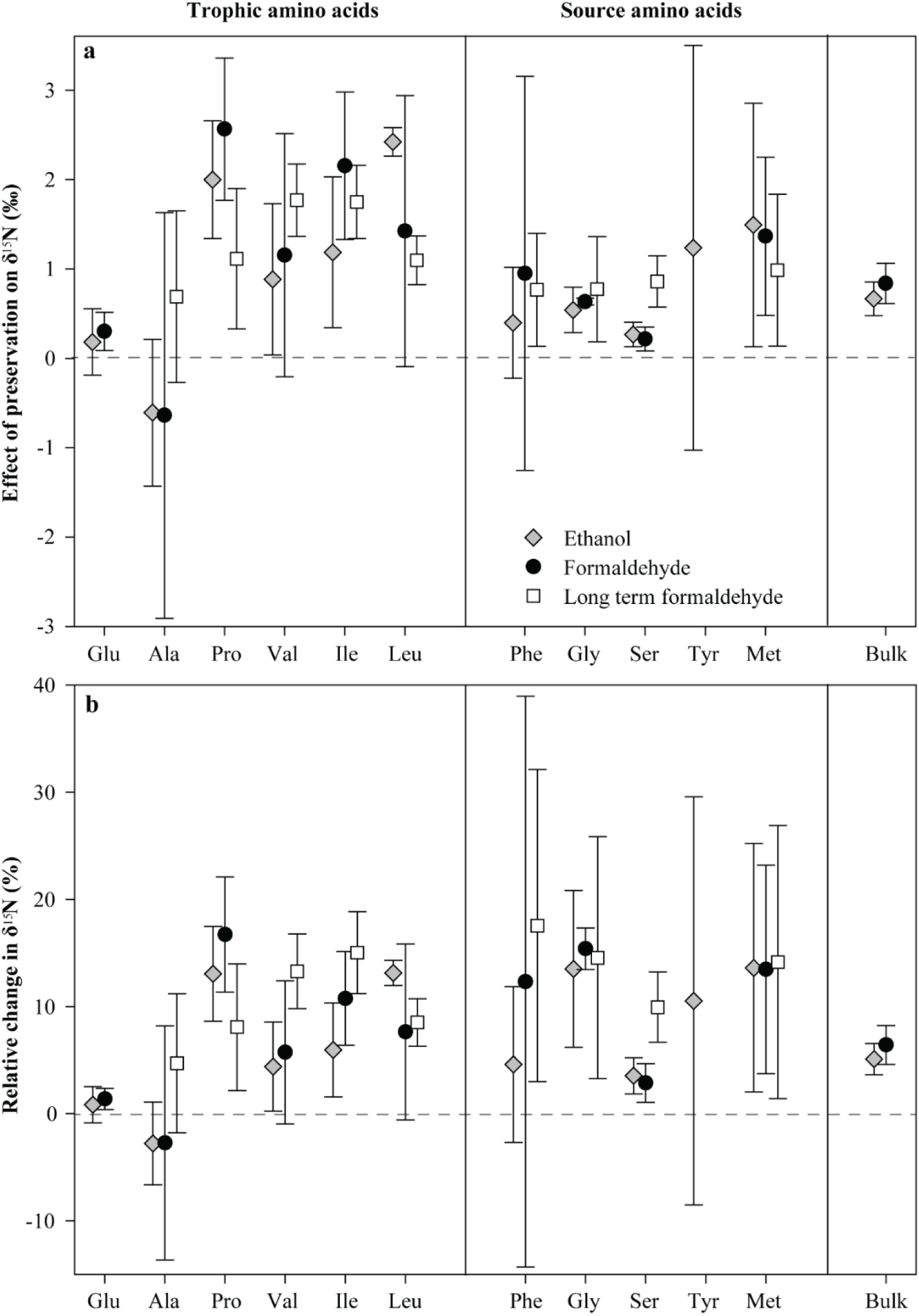
δ^15^N in short-term (1½ year) ethanol and formaldehyde preserved adult anchovy bulk material (n = 6 per treatment) and in individual AAs (Glu, Ala, Pro, Phe, Gly, n = 4; Val, Ile, Leu, Ser, Tyr, n = 3; Met, n = 2), and from long-term (27 year, n = 3 per treatment) formaldehyde-preserved larval anchovy after subtracting frozen control samples. Changes relative to controls are shown in a) ‰ and as b) relative change in δ^15^N (±1 SD). Glu could not be purified from larval anchovy, and Tyr from formaldehyde-preserved samples. n = minimum number of replicates per treatment.

Overall, short-term chemical preservation resulted in significant δ^15^N_AA_ fractionation relative to frozen control samples (F = 15.44, df = 2, p < 0.001), but did not differ between ethanol and formaldehyde (p = 0.410). Some degree of δ^15^N enrichment was observed for all short-term preserved AAs, with the exception of Ala, which was depleted. Pairwise testing revealed that Pro (+2.00 ‰, t = 6.07, df = 3, p = 0.027) and Leu (+2.42 ‰, t = 26.44, df = 2, p < 0.001) changed significantly in ethanol preserved samples. Substantial δ^15^N enrichment of 0.9 ‰ or more was also observed for Val, Ile, Tyr and Met, but with some variability between replicate measurements (Fig. 4-a, Table S1). Similarly, δ^15^N enrichment was also generally observed for all short-term formaldehyde preserved AAs, with Pro (+2.56 ‰, t = 6.45, df = 3, p = 0.023) and Gly (+0.63 ‰, t = 32.92, df = 3, p < 0.001) changing significantly. Enrichment of 0.9 ‰ or more was observed for Val, Leu, Phe and Met, but varied between replicate measurements (Fig. 4-a, Table S1). Glu and Ser were the least affected by short-term ethanol and formaldehyde preservation (+0.3 ‰ or less).

Long-term formaldehyde preservation of small 8.5-10 mm SL anchovy larvae also resulted in fractionation of δ ^15^N_AA_ (F = 80.58, df = 1, p < 0.001). Fractionation patterns differed slightly, but not significantly from those observed for short-term formaldehyde preserved adult anchovy (Fig. 4-a, Table S1, F = 0.02, df = 1, p = 0.904). Here, significant effects were seen for Val (+1.77 ‰, t = 7.61, df = 2, p < 0.017), Ile (+1.75 ‰, t = 7.41, df = 2, p < 0.018), Leu (+1.10 ‰, t = 6.96, df = 2, p = 0.020) and Ser (+0.86 ‰, t = 5.21, df = 2, p < 0.035). Least affected by long-term formaldehyde preservation were Ala, Phe and Gly (+0.7-0.8 ‰).

We observed similar patterns of preservation effects across a suite of AAs irrespective of the preservative used. Although formaldehyde preservation caused added variability (particularly for Ala, Val, Leu and Phe), there were no differences overall in average variabilities between control samples and any of the treatments (Supplementary Table S1). The slightly lower variability for long-term formaldehyde-preserved anchovy may have been the result of 6-11 larvae being pooled together per sample. Relative δ^15^N change was more consistent among AAs (Fig. 4-b) than absolute change, which was generally higher for trophic compared to source AAs (Fig. 4-a). Unfortunately, it was not possibly to purify Glu from the anchovy larvae samples. In addition, Tyr was only be detected in frozen and ethanol-preserved samples.

### Consequences for trophic position estimates

To evaluate the use of preserved specimens for calculating TPs of fish, we compared different estimates derived from conventional trophic and source AA combinations (Fig. 5). Due to substantial fractionation of Ile, Met and Tyr during purification, we did not use these AAs in our TP estimates. We found no effect of preservative or preservative:AA interaction on TP estimates for short-term formaldehyde (F = 0.64, df = 1, p = 0.428; F = 1.68, df = 14, p = 0.080) or ethanol-preserved (F = 3.34, df = 1, p = 0.072; F = 1.56, df = 14, p = 0.113) samples compared to frozen samples. The same was seen for long-term formaldehyde preserved samples (F = 3.94, df = 1, p = 0.053; F = 0.44, df = 11, p = 0.930). However, low p values do point to a trend in changing TP estimates as illustrated in Fig 5. Furthermore, this TP difference varied depending on the combination of trophic and source AAs.

**Figure 5:**
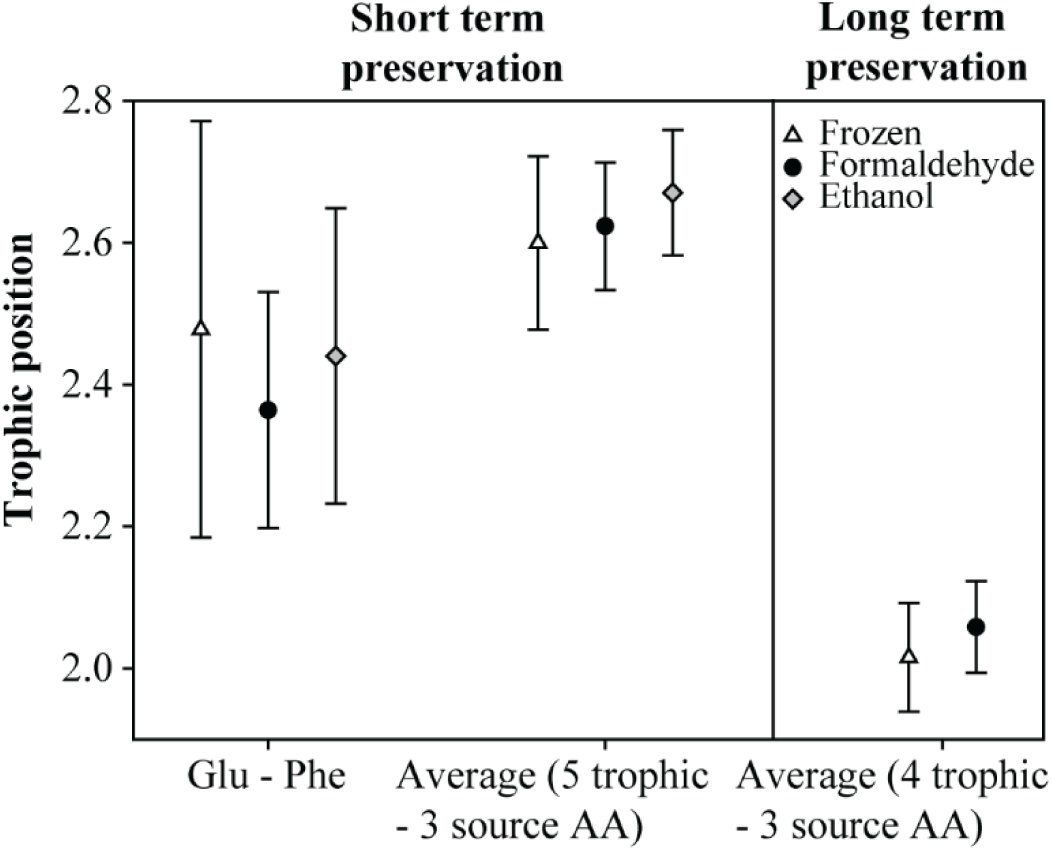
Comparison of TP estimates of frozen, ethanol and formaldehyde preserved samples calculated from Glu and Phe (n = 4) and as an average of all TP estimates (n = 3) based on Glu, Ala, Pro, Val, Leu, Phe, Gly and Ser (±SD). Glu could not be purified from the long-term preserved larval anchovy.

## Discussion

Here we demonstrate that chemical preservation can cause statistically significant isotopic fractionation of N in some ecologically relevant AAs but that 27-year long-term preservation did not result in additional fractionation. These results were achieved using a novel CSIA-AA approach with a high analytical precision. Despite the observed fractionations, N source and TP estimates were minimally affected by preservation, particularly when using multiple trophic and source AAs. Nevertheless, it is important to consider ecological and biogeochemical ramifications of chemical preservation, as well as methodological uncertainty in experimental design.

### Performance of the HPLC/EA-IRMS approach

The High Pressure Liquid Chromatography Elemental Analysis – Isotope Ratio Mass Spectrometry (HPLC/EA-IRMS) method successfully provided δ^15^N measurements across a suite of standard amino acids (AAs). Sample processing and purification by HPLC resulted in low isotopic fractionation of nitrogen (N) for most AAs concerned. Glu, Ala and Phe, particularly important AAs in ecological studies (e.g., Chikaraishi et al. 2009; McMahon and McCarthy 2016; Décima et al. 2017), demonstrated almost no fractionation relative to control samples and little variation among replicate measurements. Broek and McCarthy (2014) reported a mean accuracy of 0.12 ± 0.05 ‰ using Glu and Phe powdered standards, slightly below the 0.02 ± 0.02 ‰ that we observed for the same AAs. The improved accuracy in our study may have resulted from pre-purification of our Pierce AA standard mix, a decision made after observing high fractionation in Phe powdered standards during preliminary testing, which suggested N containing contaminants in the standard. Overall, we achieved high accuracy for most of our AA standards (0.06 ± 0.21 ‰, Fig. 3) except for Ile, Tyr and Met, which underwent substantial fractionation. Tyr and Met are particularly prone to oxidation (e.g., Sprung et al. 2009), which could explain the high δ^15^N enrichment. These findings are largely in agreement with those of Broek et al. (2013), who also observed high fractionation for Ile and Met.

Interestingly, our procedural reproducibility error was equal to the precision of the IRMS instrument, meaning that neither sample handling during the multiple processing steps nor purification by HPLC added any quantifiable variability to the analysis (Fig. 3). By comparison, Broek and McCarthy (2014) observed a precision of 0.3 ‰ for Glu and Phe, which was half that of their control samples. However, it is important to note that all these tests are based on AA standards rather than organism samples. Thus, the samples did not undergo hydrolysis, a step that may impact accuracy and procedural reproducibility. Using Phe from a hydrolyzed cyanobacteria culture, Broek et al. (2013) observed a larger procedural reproducibility error (0.55 ‰) than when using Phe standards. The complex molecular structure of organismal samples relative to laboratory AA standards means that other N-containing compounds may co-elute with collected AAs adding additional analytical variability. Indeed, co-elution with an unknown compound was the reason Glu could not be purified in some of our larval anchovy samples, although, for most anchovy AAs concerned, the chromatograms showed no co-elution with unknown compounds (Supplementary Fig. S1).

The HPLC/EA-IRMS method performed well relative to the conventional GC-C-IRMS method generally used for CSIA-AA. Recent aquatic studies using GC-C-IRMS report average precisions in the range of 0.4 – 1.0 ‰, but with considerable variability among analyses (Broek et al. 2013; Ogawa et al. 2013; Broek and McCarthy 2014; Chikaraishi et al. 2015; e.g., Bradley et al. 2016; Hetherington et al. 2017; Ruiz-Cooley et al. 2017; Nuche-Pascual et al. 2018; Vane et al. 2018; Vokhshoori et al. 2019). In the literature, such error reporting is often based on only one to a few AA standards (Broek et al. 2013), which for the most part have not gone through any of the sample processing steps prior to derivatization. In addition, errors are often expressed as the standard deviation of replicate injections of the same derivatized standard or biological sample, thus giving only the analytical precision of the instrument, not procedural reproducibility. Nonetheless, these published errors are more than double our procedural reproducibility of 0.15 ± 0.07 ‰, and the reproducibility error of the GC-C-IRMS method is likely to be higher, e.g., from slight inconsistencies in the derivatization procedure (see Broek et al. 2013; Broek and McCarthy 2014). Derivatization of AAs can also cause significant fractionation (e.g., by 2.5 ± 1.2 ‰ in Broek and McCarthy 2014), with δ^15^N values typically adjusted following correction protocols. Using the HPLC/EA-IRMS method, we found no significant fractionation in 8 of the 11 AAs tested, thus eliminating the need for correcting these 8 AAs. If Ile, Tyr and Met are used, however, we do advise adjusting their δ^15^Ns by the observed change as described in our assessment.

### Effects of chemical preservation and storage time

The preservation of adult anchovy white muscle in ethanol or formaldehyde caused significant fractionation of δ^15^N compared to frozen samples (+0.67 and +0.84 ‰, respectively). We used frozen samples as controls since freezing has been shown not to cause N fractionation in fish and is generally considered the safest method of preservation (Bosley and Wainright 1999; Kaehler and Pakhomov 2001; Sweeting et al. 2004). Past studies with a variety of aquatic organisms have found inconsistent effects of preservatives (e.g., Kelly et al. 2006; Barrow et al. 2008). However, our findings are within the range of published values (+0.6 to +1.4 ‰) for multiple species of marine fin fish (Bosley and Wainright 1999; Kaehler and Pakhomov 2001; Arrington and Winemiller 2002; Sweeting et al. 2004; Kelly et al. 2006; Hetherington et al. 2019).

We also found significant preservation effects for several individual AAs. Notably, Pro and Leu showed changes in δ^15^N irrespective of the preservative used, and Ile, Val, Gly and Ser also changed in the formaldehyde treatments. This is not entirely surprising given the observed isotopic changes in bulk N, which is mainly stored in AAs. Nevertheless, of the five other published studies of preservative effects on δ^15^N in individual AAs, ours is the first to observe significant differences. For several species of freshwater fishes, Ogawa et al. (2013) and Chua et al. (2020) concluded that formaldehyde preservation had no effect. Similarly, Hetherington et al. (2019) did not find significant changes for yellowfin tuna AAs preserved in ethanol and formaldehyde despite observing significant changes in bulk values. Studies on zooplankton, squid tissue and corals have also reported no significant change (Hannides et al. 2009; Hetherington et al. 2019; Strzepek et al. 2014). Although a comparison of these studies point potentially toward species-specific differences in preservative effects, another reason why the previous studies did not find significant effects is likely due to the choice of CSIA-AA method combined with low number of replicates. Since the GC-C-IRMS method used in these previous studies has an average precision of 0.5-1.0 ‰ (Hannides et al. 2009; Ogawa et al. 2013; Strzepek et al. 2014; Hetherington et al. 2019; Chua et al. 2020), it cannot resolve isotopic preservation effects of comparable 1 ‰ magnitude (this study; Hetherington et al. 2019).

How nitrogen is fractionated during sample preservation is not clear, although it appears to be selective for specific AAs rather than all AAs changing in proportion to bulk N isotopes. Neither ethanol nor formaldehyde contains any N that could be added to the bulk tissues or individual AAs thereby altering the N isotopic ratios. Although not expected, if proteins are partially hydrolyzed during preservation resulting in C-^14^N bonds being preferentially cleaved, this could result in N leakage into the preservative solution as, e.g., free AAs of amines (Silfer et al. 1992; Bosley and Wainright 1999; Sarakinos et al. 2002). In bulk tissue samples, we did register a modest reduction in N content when preserved in formaldehyde (−1.5 ± 0.9 %), but for ethanol the N content increased (+1.5 ± 0.9 %) relative to frozen samples. Ethanol is an organic solvent that is well suited for extracting lipids from tissues (Kelly et al. 2006), which may have masked any N loss in the ethanol-preserved samples. This does not explain, however, why some AAs appear to fractionate more than others. Hetherington et al. (2019) tested whether there was a greater loss of high compared to low fractionating AAs using peak areas, but found no differences. For our analyses, however, the two AAs (Pro and Gly) that fractionated significantly in the ethanol- and formaldehyde-preserved adult anchovy (Fig. 4) showed significantly greater loss of mass relative to frozen control samples than did the two lowest fractionating AAs (Glu and Ser, p ≤ 0.05, supplementary Table S2). Future studies addressing the effects of chemical preservatives should attempt to determine which compounds are lost from the tissues and end up in the preservative solution. It may also be informative to test how preservatives impact the hydrolysis of different proteins and the acid lability of certain amine bonds that could effect hydrolysis and subsequent recovery of specific AAs.

Another possible reason for the fractionation in preserved samples is co-elution with N-containing compounds during purification by HPLC. Indeed, we did see an unknown compound co-eluted with Ala in frozen adult anchovy samples, which may explain why Ala was the only AA depleted in δ^15^N in the preserved samples (Supplementary Fig. S1-a). δ^15^N gradients have been observed across chromatographic peaks of eluting compounds, with the tail ends being considerably enriched (Hare et al. 1991; Broek et al. 2013). During collection, part of the front end of the Ala peak was mixed with the tail end of the unknown compound. Assuming this compound contained N, it may well have elevated the δ^15^N in our control samples. Broek and McCarthy (2014) also observed that amino sugars elute just prior to Glu, which was the reason we did not attempt to collect Glu from the frozen larval anchovy samples. However, for most other AAs, there was no indication of co-elution, indicating that poor chromatography could not have driven the general fractionation pattern that we observed (Supplementary Fig. S1). In fact, we actually observed improved chromatographic separations of AAs in both the ethanol and formaldehyde preserved samples. Hetherington et al. (2019) also suggested that co-elution was responsible for some of the observed δ^15^N_AA_ fractionation, and that chemical preservation may have aided purification and improved their gas chromatography.

Overall, the effects of ethanol and formaldehyde preservation on δ^15^N_AA_ were quite similar and did not increase variability in replicate measurements. Preserving anchovy in formaldehyde for 1½ or 27 years also did not significantly alter the fractionation patterns. These results are promising for comparing different samples in historical archives, where the duration of preservation can vary considerably and where different preservatives are often used. However, it is important to differentiate between statistically quantifiable effects and ecologically or biogeochemically relevant offsets. The substantial, and in some cases significant, enrichment in δ^15^N_AA_ does illustrate that caution must be used in interpreting results and that it may be necessary to apply corrections, depending on the questions, organisms, systems being studied. This would be particularly important where the aim is to combine or compare preserved with fresh or frozen samples, or when it is necessary to know the true δ^15^N value. Natural variability in the δ^15^N of different N sources or across isoscapes is often on the order of a few permille (e.g., Ohkouchi et al. 2017). Thus, studies attempting to assess the relative contributions of N sources to the food chains of target organisms by comparing source AA Phe δ^15^N signatures with inorganic N isotopes in the environment should correct Phe for formaldehyde and ethanol preservation effects (−0.9 and −0.4 ‰, respectively; see supplementary Table S1). Considering the uneven fractionation patterns across different AAs, studies that utilize isotopic fingerprinting or mixing models to determine dietary contributions of different prey may also consider correcting for preservation effects. Such correction factors may need to be species-specific (e.g., Kelly et al. 2006), given the large δ^15^N_AA_ fractionation differences of various organisms (this study; Hetherington et al. 2019). Although preservation did not result in large shifts in TP estimates, the up to 0.12 TP difference between preserved samples can be ecologically significant (Fig. 5). For instance, assuming a 10 % trophic transfer efficiency, the formaldehyde preserved TP estimate (Glu-Phe) would overestimate energy transfer from the base of the food web to adult anchovy by ∼12 %. For studies aimed at quantifying mass balances using stable isotopes, it may also be necessary to correct individual AAs by the observed fractionation (supplementary Table S1). Finally, variable δ^15^N_AA_ fractionation highlights the need for careful consideration in the choice of AAs to analyze. Nielsen et al. (2015), amongst others, have advocated for the use of multiple trophic-source AA combinations when calculating TPs. In light of our results, this may be particularly important when working with preserved samples.

### Comments and recommendations

This study contributes to the growing understanding of the effects of preservatives on individual AAs. For Northern Anchovy, we show that the duration of chemical preservation and the preservative used (ethanol or formaldehyde) in an historical archive results in only a small alteration of δ^15^N_AA_. Compared to non-chemical preservation, significant δ^15^N_AA_ alteration can be expected and may, to some extent, be species or case specific. We recommend that studies involving different species or preservatives should consider short-term pilot testing to assess the need for correction protocols. Nonetheless, our results are also encouraging for reconstructing biogeochemical records and food web trophic connections over long time scales. A vast amount of the information locked away in historical archives of preserved samples can now be accessed using CSIA-AA.

This study has also helped demonstrate the possibilities of an HPLC/EA-IRMS approach for CSIA-AA. The superior precision and accuracy of this method are particularly suitable for studies attempting to resolve fine-scale variability in N sources and TPs. Among primary or secondary consumers in marine pelagic systems, for instance, temporal and spatial variability in TP is usually on the order of ±0.4 TP (Hannides et al. 2009; Choy et al. 2012; Décima et al. 2013; Choy et al. 2015; Miyachi et al. 2015; Bradley et al. 2016; Laiz-Carrión et al. 2019). Even small shifts in TP can be associated with significant food web disruptions and represent considerable changes in energy transfer to higher level consumers (Vander Zanden et al. 1999). The propagation of methodological errors reported in this study equals a ±0.04 methodological uncertainty in TP estimates based on Glu and Phe, the most widely used trophic-source AA combination (TDF of +5.7, Bradley et al. 2015). In comparison, the analytical errors reported for the GC-C-IRMS approach is equivalent to ±0.1 - 0.25 TP uncertainty (see discussion), within the range of natural variability for many marine organisms.

Nonetheless, before opting for the HPLC/EA-IRMS approach, one should consider a number of issues as this approach may not always be better than conventional GC-C-IRMS. First and foremost, the method is only as good as the quality of the chromatography and the precision of the IRMS. Samples with complex biochemical compositions, such as degraded organic matter, can result in messy chromatograms rendering AA purification difficult or impossible (Broek and McCarthy 2014, and references therein). Although not shown here, preliminary testing on crustacean zooplankton revealed poorer chromatographic performance than for frozen larval anchovy (Supplementary Fig. S2-d). Crustaceans notably have large amounts of polysaccharide chitin in their exoskeleton that breakdown to monosaccharides when hydrolyzed (e.g., GlcNAc and GlcN, Einbu and Vårum 2008). These monosaccharides can co-elute with Gly, Thr and Glu and prevent purification (present study; Broek and McCarthy 2014). For larval fish, which contain chitin in their epidermis (Tang et al. 2015) along with cartilage, this issue seems to abate with development and increasing size, but for small crustaceans, the HPLC/EA-IRMS method may not be appropriate. Due to the low concentrations of some AAs (such as Phe), the N collected from a single HPLC run may also not be within the detection range of most IRMS systems. Injecting too much sample onto the column reduces chromatographic performance, and combining multiple collections increases time and costs. One solution is to use high-sensitivity Nano-EA-IRMS instrumentation as was done by Broek and McCarthy (2014) and in the present study, which allows CSIA-AA on individual fish larvae of ≥350 µg DW. Lastly, although HPLC/EA-IRMS has the potential to be a relatively fast CSIA-AA method (we routinely processed 25 or more samples per week), it can still be as costly as GC-C-IRMS if a large suite of AAs is desired, since every AA needs to be analyzed for δ^15^N_AA_ individually.

## Acknowledgements

We would like to thank Taylor Broek and Matthew McCarthy for their inputs on implementation of the HPLC/EA-IRMS method and a special thanks to Dyke Andreasen for his tireless pursuit in optimizing the Nano-EA-IRMS system at the University of California, Santa Cruz at Stable Isotope Laboratory facility. We would also like to thank Magali Porrachia and Dereka Chargualaf for their assistance in the lab and Bruce Deck for his efforts in optimizing the EA-IRMS system at Scripps Institution of Oceanography. Finally, we thank William Watson and our reviewers for commenting on the manuscript. This work was supported by the Danish Council for Independent Research and the Marie Curie COFUND program (grant ID: DFF – 4090-00117 to R.S.), NOAA Fisheries and the Environment (FATE grant to A.R.T.), from NOAA RESTORE Science Program (grant ID: NA15OAR4320071 to M.R.L), and from NSF to the California Current Ecosystem LTER site.

## Supplementary material

**Table S1:** δ^15^N of bulk material and individual AAs (±1 SD, n in brackets) in short-term (∼1½ year) frozen, ethanol and formalin preserved adult anchovy white muscle tissue and long-term (27 year) frozen and formaldehyde preserved larval anchovy. na: not analyzed. nd: not detected. * denotes significant effects of chemical preservation compared to frozen samples at the 0.035 level.

| AA | Short term preservation $\delta^{15}\text{N}$ | | | Long term preservation $\delta^{15}\text{N}$ | |
| --- | --- | --- | --- | --- | --- |
|  | Frozen | Ethanol | Formaldehyde | Frozen | Formaldehyde |
| Glu | $22.2 \pm 0.26$ (4) | $22.39 \pm 0.27$ (4) | $22.51 \pm 0.18$ (4) | na | na |
| Ala | $21.05 \pm 1.01$ (4) | $20.44 \pm 0.62$ (4) | $20.41 \pm 1.65$ (4) | $15.42 \pm 0.78$ (3) | $16.11 \pm 0.19$ (3) |
| Pro | $15.37 \pm 0.25$ (4) | $17.37 \pm 0.41$ (4)* | $17.94 \pm 0.61$ (4)* | $14.04 \pm 0.52$ (3) | $15.15 \pm 0.34$ (3) |
| Val | $20.85 \pm 1.08$ (3) | $21.73 \pm 0.24$ (3) | $22.00 \pm 0.77$ (3) | $13.41 \pm 0.63$ (3) | $15.18 \pm 0.32$ (3)* |
| Ile | $20.19 \pm 0.72$ (3) | $21.38 \pm 0.65$ (3) | $22.35 \pm 0.43$ (3) | $11.68 \pm 0.26$ (3) | $13.43 \pm 0.26$ (3)* |
| Leu | $18.45 \pm 0.51$ (3) | $20.87 \pm 0.39$ (3)* | $19.87 \pm 1.87$ (3) | $12.91 \pm 0.15$ (3) | $14.01 \pm 0.14$ (3)* |
| Phe | $10.18 \pm 1.62$ (4) | $10.58 \pm 1.29$ (4) | $11.13 \pm 1.01$ (4) | $4.49 \pm 0.23$ (3) | $5.26 \pm 0.41$ (3) |
| Gly | $4.15 \pm 0.41$ (4) | $4.69 \pm 0.20$ (4) | $4.79 \pm 0.39$ (4)* | $5.42 \pm 0.64$ (3) | $6.19 \pm 0.76$ (3) |
| Ser | $7.53 \pm 0.40$ (3) | $7.80 \pm 0.49$ (3) | $7.74 \pm 0.43$ (3) | $8.65 \pm 0.08$ (3) | $9.51 \pm 0.31$ (3)* |
| Tyr | $13.37 \pm 1.36$ (3) | $14.60 \pm 0.96$ (3) | nd | na | nd |
| Met | $10.52 \pm 1.04$ (2) | $12.01 \pm 2.4$ (2) | $11.89 \pm 0.15$ (2) | $7.35 \pm 0.68$ (3) | $8.34 \pm 0.33$ (3) |
| Bulk | $13.12 \pm 0.25$ (6) | $13.79 \pm 0.29$ (6)* | $13.96 \pm 0.15$ (6)* | na | na |

**Table S2:** Mass of high-fractionating (Pro, Gly) relative to low-fractionating (Glu, Ser) AAs in adult anchovy white muscle tissue (±1 SD). * indicate test samples that differ significantly from control samples at the 0.05 level (AVOVA).

| AA ratio | Control<br>Frozen | Test |  |
| --- | --- | --- | --- |
|  |  | Formaldehyde | Ethanol |
| Pro:Glu | $0.32 \pm 0.01$ | $0.32 \pm 0.02$ | $0.29 \pm 0.02^*$ |
| Gly:Glu | $0.36 \pm 0.01$ | $0.33 \pm 0.02$ | $0.31 \pm 0.03^*$ |
| Pro:Ser | $1.19 \pm 0.05$ | $0.99 \pm 0.05^*$ | $0.95 \pm 0.16^*$ |
| Gly:Ser | $1.33 \pm 0.06$ | $1.02 \pm 0.06^*$ | $1.05 \pm 0.20^*$ |

**Figure S1:**
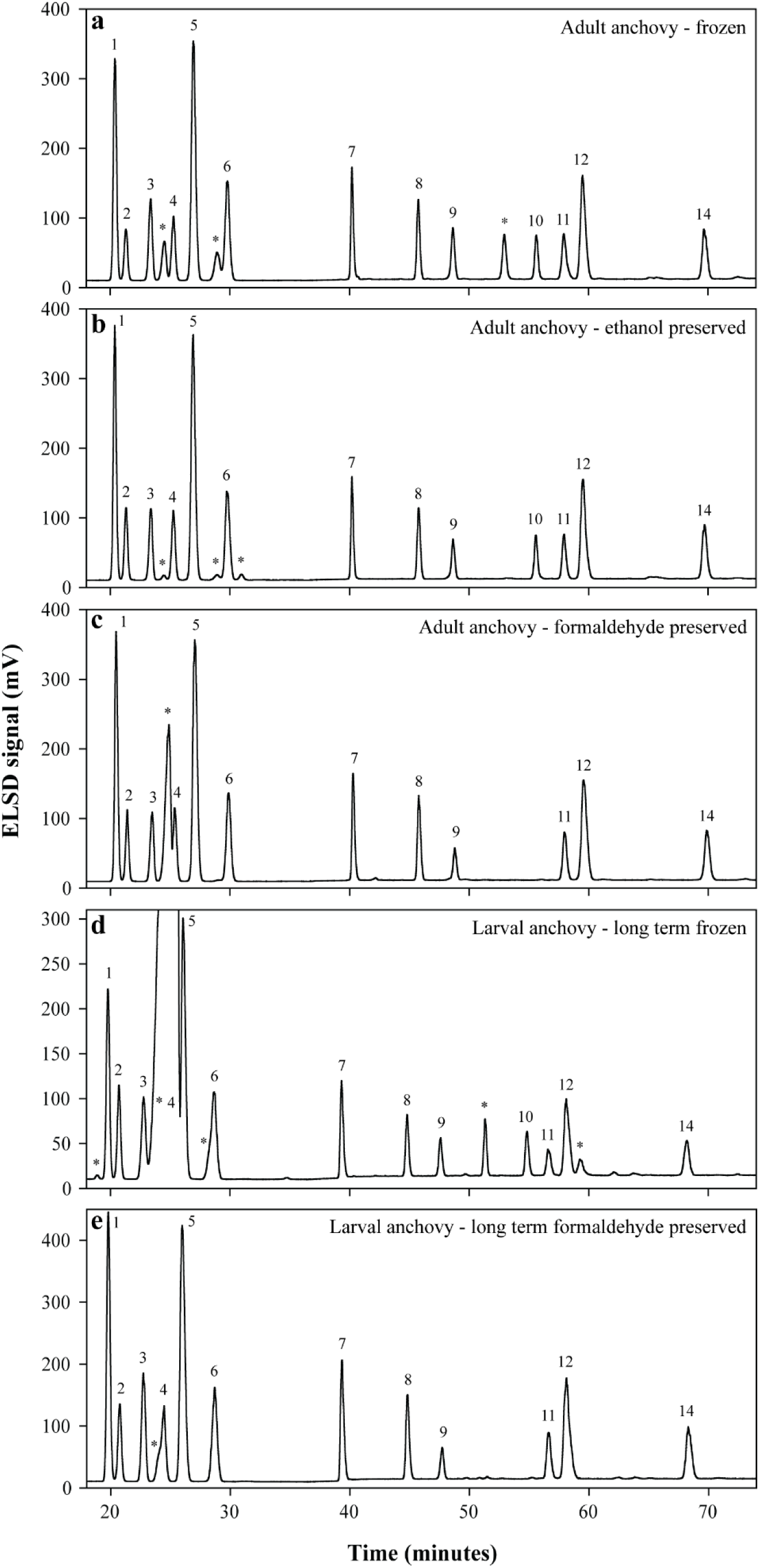
Chromatograms from representative samples of adult anchovy white muscle tissue preserved a) frozen at −20°C, b) in ethanol and c) in formaldehyde for ∼1½ years, and pooled samples of 8.5-10 mm SL sized larval anchovy preserved d) frozen at −80°C and e) in formalin for 27 years. Peak identities are: 1. Asp, 2. Ser, 3. Gly, 4. Thr, 5. Glu, 6. Ala, 7. Pro, 8. Val, 9. Met, 10. Tyr, 11. Ile, 12. Leu, 13. Cys, 14. Phe. * marks unknown compounds. Purification of Gly, Glu and Ala was best achieved in formaldehyde preserved samples. Slight co-elution of Ala and unknown compounds occurred in frozen samples and for Gly in larval frozen samples. Glu could not be purified in larval frozen samples, and Tyr was undetectable in formaldehyde preserved samples. Note that the amount of sample mass injected varied between runs on the HPLC.

